# Quantifying rhythmicity in perceptual reports

**DOI:** 10.1101/2022.05.16.492063

**Authors:** Tommaso Tosato, Gustavo Rohenkohl, Jarrod Robert Dowdall, Pascal Fries

## Abstract

Several recent studies investigated the rhythmic nature of cognitive processes that lead to perception and behavioral report. These studies used different methods, and there has not yet been an agreement on a general standard. Here, we present a way to test and quantitatively compare these methods. We simulated behavioral data from a typical experiment and analyzed these data with several methods. We applied the main methods found in the literature, namely sine-wave fitting, the Discrete Fourier Transform (DFT) and the Least Square Spectrum (LSS). DFT and LSS can be applied both on the averaged accuracy time course and on single trials. LSS is mathematically equivalent to DFT in the case of regular, but not irregular sampling - which is more common. LSS additionally offers the possibility to take into account a weighting factor which affects the strength of the rhythm, such as arousal. Statistical inferences were done either on the investigated sample (fixed-effect) or on the population (random-effect) of simulated participants. Multiple comparisons across frequencies were corrected using False-Discovery-Rate, Bonferroni, or the Max-Based approach. To perform a quantitative comparison, we calculated Sensitivity, Specificity and D-prime of the investigated analysis methods and statistical approaches. Within the investigated parameter range, single-trial methods had higher sensitivity and D-prime than the methods based on the averaged-accuracy-time-course. This effect was further increased for a simulated rhythm of higher frequency. If an additional (observable) factor influenced detection performance, adding this factor as weight in the LSS further improved Sensitivity and D-prime. For multiple comparison correction, the Max-Based approach provided the highest Specificity and D-prime, closely followed by the Bonferroni approach. Given a fixed total amount of trials, the random-effect approach had higher D-prime when trials were distributed over a larger number of participants, even though this gave less trials per participant. Finally, we present the idea of using a dampened sinusoidal oscillator instead of a simple sinusoidal function, to further improve the fit to behavioral rhythmicity observed after a reset event.

## 1. Introduction

Brain activity typically shows distinct rhythms, which entail fluctuations in excitability of groups of neurons located in specific areas. The effect of such rhythms on behavior can be tested using the appropriate experimental design. There are at least two different approaches allowing to do so. The first one consists in showing a dependence of behavior on the phase of a neuronal rhythm measured by EEG or MEG. This has been done for theta (Busch et al., 2009; Landau et al., 2015; Wutz et al., 2016), alpha (Guo et al., 2014), beta (Baumgarten et al., 2015), and gamma (Ni et al., 2016). The second approach consists in showing rhythmicity directly on behavior, aligning task performances to an event, which can be internally or externally generated. This event is usually paired to a detection or discrimination task, with the probe being presented at variable time intervals from the reset event. Externally generated events consist of auditory stimulations (Dehaene, 1993; Romei et al., 2012), visual stimulations (Landau and Fries, 2012; Fiebelkorn et al., 2013), or TMS pulses (Veniero et al., 2021). Externally generated events may act as an alignment event by resetting the phase of internal rhythms as they directly interfere with the neural activity in the respective sensory areas, or by resetting attentional dynamics. Conversely, examples of internally generated events are motor acts like an arm movement (Tomassini et al., 2015), a button press (Benedetto et al., 2016) or an eye movement (Bellet et al., 2017; Benedetto and Morrone, 2017). Internally generated events may also act as an alignment event by resetting neural rhythms, either through a corollary discharge, i.e. an efferent copy of the movement plan sent by motor areas, or through the generation of new sensory inputs, e.g. the retinal movement during a saccade. Alternatively, or additionally, a motor action may act as an alignment event by revealing an internal rhythm, if it is produced with some preference for a particular phase of that rhythm.

When the rhythmicity of behavioral metrics is directly quantified, there are several challenges. First, the data are very sparse: each trial provides only one measure of behavioral performance (e.g. hit or miss), which alone does not provide any information about rhythmicity. Second, the sampling of the data can be irregular, and this can be a challenge for traditional frequency analysis methods such as Discrete Fourier Transform (DFT). Third, in the existing publications, a variety of methods for spectral analysis and for statistical testing have been used, and there is no agreement on which one offers better sensitivity and specificity.

Here we will directly compare different methods for spectral analysis and different statistical approaches, including an assessment of their strengths and weaknesses. We first generated data through a model resembling a typical experiment, we then quantified rhythmicity by various methods and we evaluated the sensitivity, specificity, and D-prime of each method.

## 2. Methods and Results

We present here the methods and results for quantifying rhythmicity in the accuracy of behavioral responses. We use the term rhythmicity to refer to the dependence of behavioral responses on the phase of a particular frequency or frequency band – but see Discussion for an elaboration on this topic. The code can be obtained here: https://github.com/tosatot/quantifying-rhythmicity-in-perceptual-reports

### 2.1. Data Generation

We simulated behavioral responses in a detection task (Fig. 1). We assumed that participants’ detection threshold is influenced by an internal rhythm, and that this rhythm is aligned across trials to an event happening at time zero. The rhythm’s modulation frequency, its modulation phase relative to the alignment event, and its modulation-depth vary somewhat across trials and participants, yet they are sufficiently consistent to result in a rhythmic modulation of the mean detection performance (Fig. 1A-C). Additionally, we simulated a factor which influences the strength of the rhythm across time. In an actual experiment, such a factor needs to be observable, e.g. pupil size, heart rate variability, skin conductance, recent performance history, recent mean reaction time, time since beginning the experiment.

**Fig. 1.**
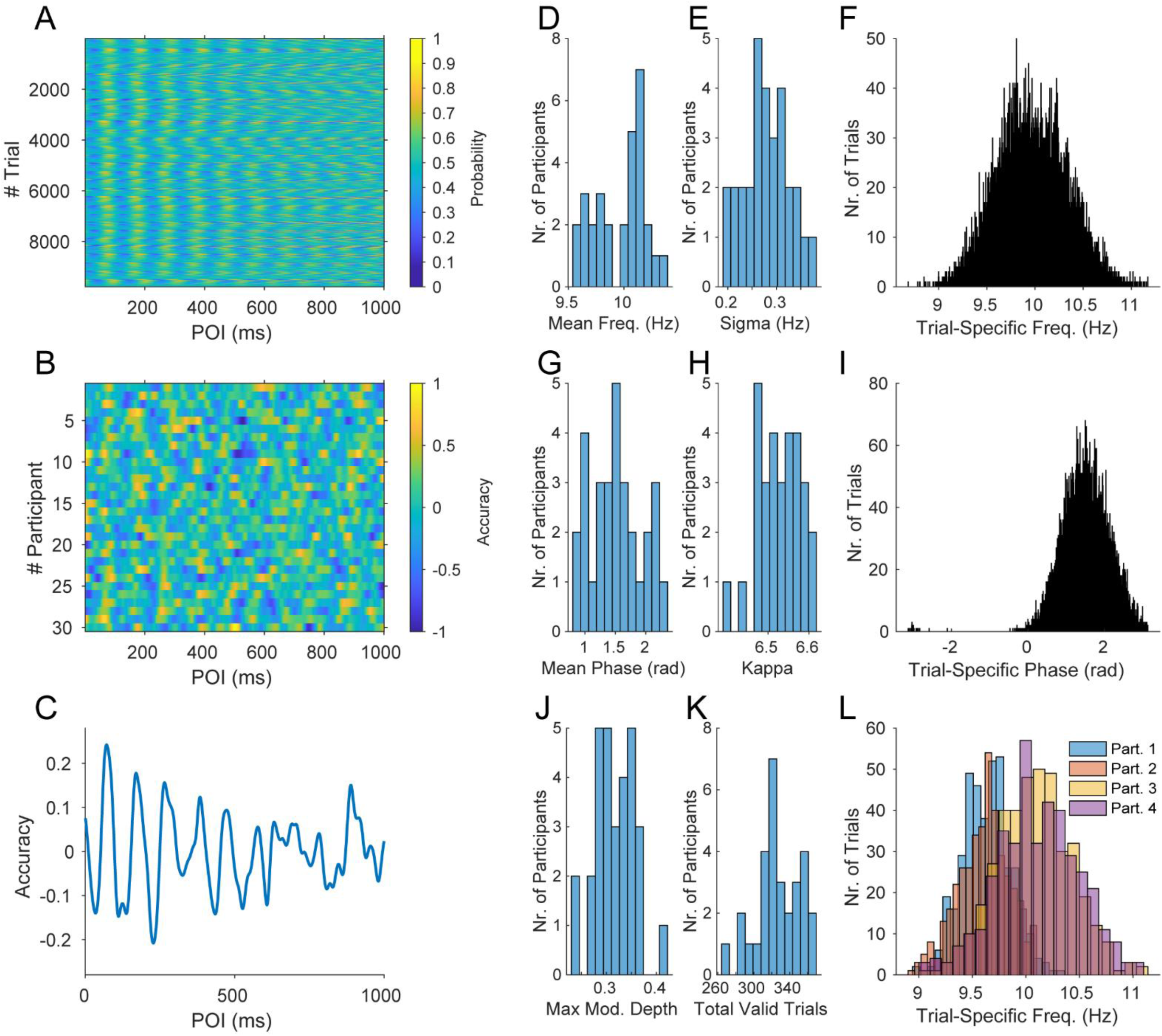
Simulation of Data. The process of generating the data is illustrated. We simulated 30 participants with 400 trials each. (A) Detection probability (color coded) as a function of probe onset interval (POI, x-axis) and trial number (y-axis, trials from all 30 participants were vertically concatenated). (B) Accuracy (color coded) as a function of POI (x-axis) and participant (y-axis). (C) Mean accuracy time course. (D) Gaussian distribution across participants of mean modulation frequencies (mean = 10 Hz, Sigma = 0.27 Hz). (E) Gaussian distribution across participants of Sigmas (mean = 0.27 Hz, Sigma = 0.05 Hz). These Sigmas determine the width of the distribution of the modulation frequency of each trial within a participant. (F) Resulting distribution across all trials of modulation frequencies. (G) Von Mises distribution across participants of mean modulation phases (mean = π⁄2 rad, Kappa = 6.53). (H) Gaussian distribution across participants of Kappas (mean = 6.53, Sigma = 0.05). These Kappas determine the width of the distribution of the modulation phase of each trial within a participant. (I) Resulting distribution across all trials of modulation phases. (J) Gaussian distribution across participants of max modulation-depth (mean = 0.32, Sigma = 0.04). (K) Gaussian distribution across participants of total valid trials (mean = 0.8, Sigma = 0.07). (L) Distribution of the modulation frequencies of each trial for 4 example participants.

Specifically, we assumed the following. 1) *Within a participant, and across trials*: the rhythm’s frequency followed a normal distribution around the participant-specific mean; its phase followed a von-Mises distribution around the participant-specific mean phase; its modulation depth varied with the arousal state. We modelled the state of arousal to decrease linearly with trial number: The first trial had maximal arousal equal to one, resulting in the predefined maximal value of modulation-depth, and the last trial had an arousal equal to zero, resulting in zero modulation-depth. Note that simulated arousal could have been assigned to different trials in any arbitrary way, and this would not have affected the analysis as long as arousal was defined (or in an experiment measurable) per trial. 2) *Across participants*: the rhythm’s participant-specific mean frequency followed a normal distribution (with means distributed as shown in Fig. 1D, unless otherwise specified for specific analyses); its participant-specific mean phase followed a von-Mises distribution (with means distributed as shown in Fig. 1G, unless otherwise specified for specific analyses); its participant-specific maximal modulation-depth followed a normal distribution (with means distributed as shown in Fig. 1J, unless otherwise specified for specific analyses); *sigma*, the width of the Gaussian distributions of trial-specific frequencies, and *kappa*, the width of the von-Mises distributions of trial-specific phases, follow a normal distributions (as shown in Fig. 1E and H, respectively); the proportion of valid trials in each participant follows a normal distribution (as shown in Fig. 1K).

We initiated the random-number generator with a unique seed for each simulation run, and we simulated each trial of each participant individually. We started by constructing the underlying probability function of each trial, which expresses the rhythmically modulated likelihood to detect a probe across different times after the aligning event. We obtained the trial-specific target frequency and the target phase randomly drawing from the frequency and phase distributions (distributions shown in Fig. 1F and I). The target frequency was used to create a third order Butterworth bandpass filter with a passband of ±0.2 Hz around the target frequency. Finally, the underlying probability function was generated by filtering white noise (i.e., uncorrelated samples uniformly distributed between 0 and 1). We discarded sufficient time (30 s) at the beginning of the signal to allow the filter to settle to its steady state. After filtering the resulting signal was visibly oscillatory, but not perfectly regular like a sine wave, because its phase did not progress linearly and its amplitude was not constant. The Hilbert transform was used to obtain the instantaneous phase, and the start time was chosen by selecting the sample that was closest to the target phase for that trial. From that moment onward, we extracted 1000 samples, corresponding to 1 s, the total duration of a trial. The amplitude of the signal was scaled between −1 to 1, and multiplied with the trial-specific modulation-depth value (obtained by randomly drawing from the modulation-depth distribution) plus 0.5. This resulted in an oscillatory signal that fluctuated around 0.5 with extrema approaching 0.5 plus/minus the modulation-depth value. This signal then represented the underlying probability function of detection performance for that trial.

We call probe onset interval (POI) the time between the alignment event and the probe event, with the latter being the stimulus event that participants are supposed to detect and report. The POI corresponds to the stimulus onset asynchrony (SOA) of several previous studies. For each simulated trial of each participant, the POI was determined as a random time between the alignment event and the end of the trial, by drawing it from a uniform distribution of values ranging from 0.001 s to 1 s in steps of 0.001 s. Note that in this process we generate an irregular sampling, e.g. not all POIs are sampled the same number of times, and some POIs do not occur at all.

Once a POI had been chosen for a trial, we looked up the value of that trial’s probability function at that POI time. We used the obtained value as probability in the following Matlab function call: “*y = datasample([-1,1],1,”weights”,[1-probability, probability])“*. Each such call gave a value of −1 or 1, with the respective probabilities summing to one. We refer to these outcome options as behavioral response value (BRV) and we consider a value of 1 to correspond to a correct detection, and a value of −1 to correspond to a missed detection.

We simulated 400 trials per participant (except otherwise noted). Only a subset of the trials were labeled as “valid”, as it is often the case in a real experiment. In each participant, trials had a certain random probability to be labeled as “valid”. This participant-specific probability was determined by randomly sampling a Gaussian distribution of mean 0.8 and sigma 0.07 (as shown in Fig. 1J).

### 2.2. Methods Evaluated

The analysis of rhythmicity, both for the simulated and any empirical data, starts with an array, which contains for each trial one POI and one corresponding BRV. In the case considered here, the POI assumes values between 0 s and 1 s, and the BRV assumes values of 1 for a correct detection, and of −1 for a missed detection.

The working hypothesis is that there is a rhythm, aligned to or reset by an event, which modulates detection performance according to the rhythm’s phase and amplitude at the moment of a probe event. The precise frequency of the rhythm is not known beforehand, therefore we need to test all plausible frequencies. That is, we need to perform the analysis in a spectrally resolved manner. Furthermore, neither the phase of the rhythm at the alignment event is known, nor the phase that leads to good or bad detection performance.

To test the working hypothesis, we need to relate POIs to BRVs, considering POIs in terms of spectral phases. We can proceed in two different ways. One option is to first combine all POIs and BRVs into an average accuracy time course (per participant, or even for the entire group of participants), and then to spectrally analyze it. Another option is to determine the spectral phases of all POIs (see below for further explanation on this) and relate them to their corresponding BRVs, e.g. through DFT or LSS.

Both approaches require estimating the spectrum of the BRVs as a function of their corresponding POIs. This spectral estimation involves or is equivalent to a Fourier transform, and therefore, we require the BRVs to be tapered to avoid leakage of spectral energy, in particular when zero padding or when testing for frequencies which are not an integer multiple of the Rayleigh frequency. Furthermore, the spectral estimation benefits from the data to be detrended before the Fourier transform. Empirical detection performance data after a reset often show longer-term trends, with the rhythms of interest superimposed. Therefore, we chose to perform a linear detrending (equivalent to a first-order polynomial detrending) of the performance timeseries, as we have done in a previous empirical study (Landau and Fries, 2012). Linear detrending removes offsets and linear trends, which can be large relative to rhythmic components, and it otherwise minimally affects the spectrum. Note that higher-order polynomial detrending can effectively constitute a high-pass filtering of the data, which renders the resulting spectra harder to interpret. Note also that we did not actively add any trend to our simulated data, such that detrending would not have been required. However, in order to make the presented methods and code directly applicable to empirical behavioral data, we included linear detrending also for the simulated data.

### 2.3. Operating on trial-averaged data

#### Preprocessing

We calculated for each participant an accuracy time course (ATC) by convolving the BRVs with a Gaussian of sigma=0.01 s in steps of 0.001 s. The convolution with a Gaussian kernel should be preferred to a box car kernel (which has been commonly used in the field). A convolution with a box car kernel in fact introduces larger and more complex distortions in the spectra than a convolution with a Gaussian (convolution in the time domain being equal to multiplication in the frequency domain).

Note that a convolution in the time domain is only necessary in the case of irregularly sampled data. For regularly sampled data (the time points are equally spaced, with each having at least one BRV associated to it, and actually each having the same number of BRVs associated to it), the ATC can be calculated by simply averaging all the BRVs corresponding to the same POI value. This avoids the low-pass filtering incurred by convolution, which is expected to reduce sensitivity for higher-frequency rhythms However, in practice it may be challenging to obtain regularly sampled data, because even if the participant is presented with an equal number of trials for each POI, some of those trials may not be accepted into the analyses, e.g. because of participant error in the task, or an artifact in the eye signal. Furthermore, if an action is used as alignment event, it is often not possible to ensure regular sampling: the POI cannot be determined a priori, e.g. because of the fixed refresh/update rate of visual displays.

The linear detrending mentioned above was applied to the ATC of each participant at this point, after the convolution. An alternative approach, followed in some previous empirical studies, is to pool all trials of all participants in the calculation of a single accuracy time course (the so called “aggregate observer”). This alternative approach does not allow to subtract a participant-specific linear trend.

#### Sine Fitting

The ATC of all participants were averaged giving a single accuracy time course. This was fitted with the sinusoidal function:

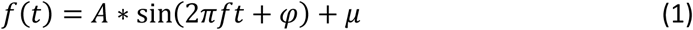

where *A, f, φ and µ* are free parameters representing respectively the amplitude, the frequency and the phase of the best sinusoidal fit, and an offset (Fig. 2A, B). This was done in Matlab using the function “*fit*” and giving “*sin1*” as the input argument for the field “*fitType*”. Statistical testing used a fixed-effect permutation approach. In each randomization, POIs and BRVs were randomly paired, and the remaining analysis was performed identically. Note that this approach results in sinewave fits with different frequencies across randomizations. Because this method finds merely the dominant spectral component, multiple-comparison correction across frequencies is not necessary. The r-squared value of the fit to the observed mean ATC was compared to the distribution of r-squared values from those randomizations, and it was considered significant if it was larger than the 95^th^ percentile (Fig. 2B). This method, with slight variation, has been used in several studies (Tomassini et al., 2015; Benedetto et al., 2016; Benedetto and Morrone, 2019).

**Fig. 2.**
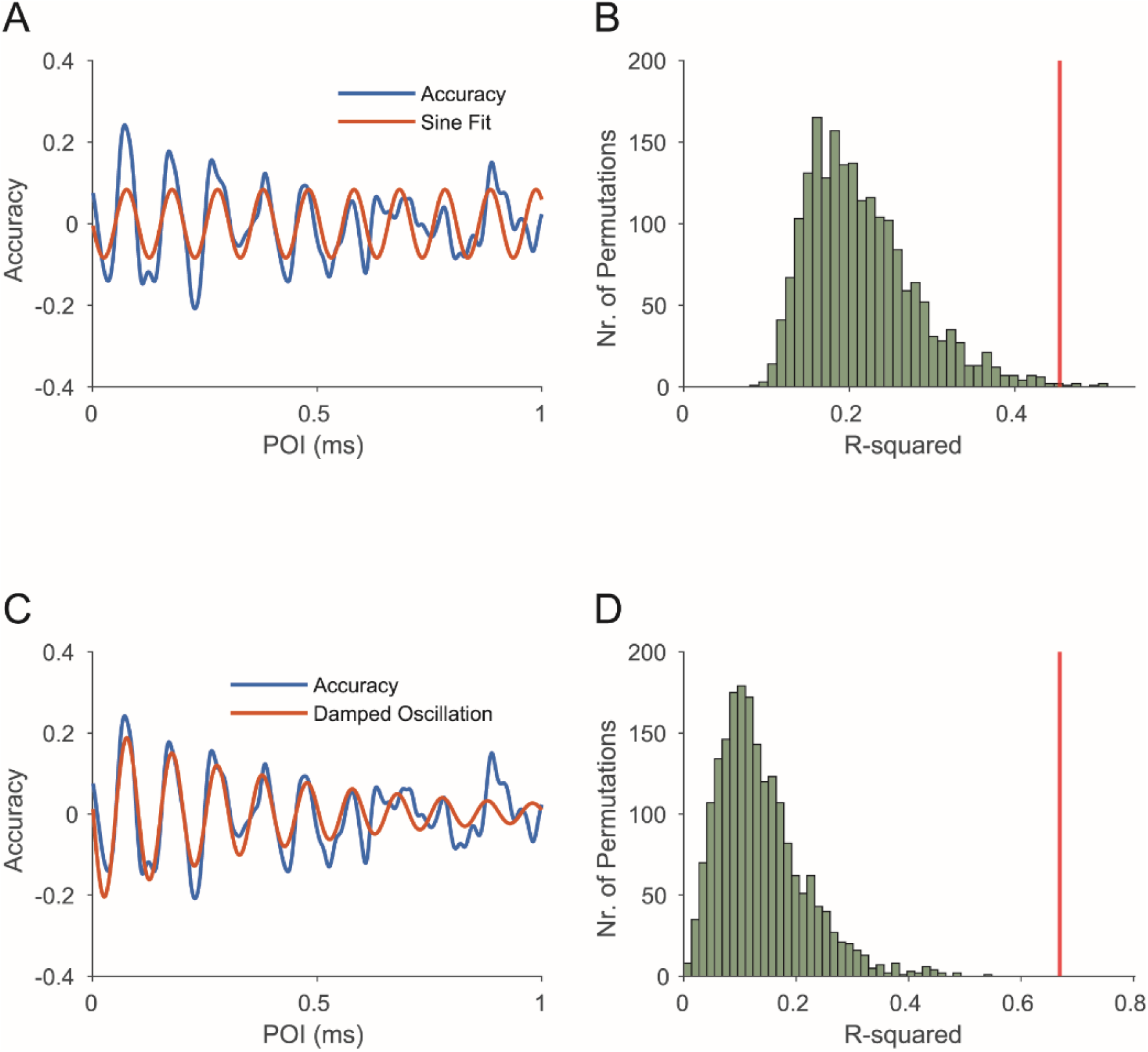
Sine Fitting. (A) Best fit of the a sine wave (red) to the mean accuracy time course (blue) generated in Fig. 1. (B) Permutation statistics to calculate the significance of the r-square value of the sine fit. The green histogram shows the r-square values of all permutations, the red line shows the r-square value of the sine fit to the observed data. (C) Best fit of a damped harmonic oscillator (red) to the mean accuracy time course (blue). (D) Permutation statistics to calculate the significance of the r-square value of the sine fit.

This approach can only provide parameters for the strongest rhythmic component, i.e. the rhythmic component explaining the largest proportion of the total variance. Thus, this approach is not suitable to capture multiple coexisting rhythms. Generally speaking, this approach does not provide a full characterization of rhythm strengths as a function of frequency, i.e. it does not provide a full spectral characterization. For this reason, we will focus the rest of this paper on methods providing full spectral characterization.

#### Accuracy Time Course Discrete Fourier Transform (atcDFT)

The ATC of each participant was first Hann tapered and zero padded to 4 seconds. We performed a discrete Fourier transform to obtain a complex spectrum per participant. The discrete Fourier transform represents a signal as the sum of a series of sinusoidal components:

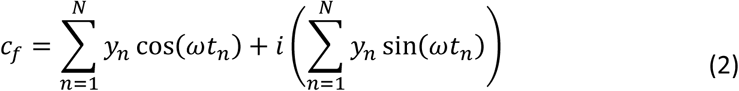

Where *y*_*n*_ is the *n*^th^ element of the vector containing the ATC, with *n* corresponding to the time bin number and *N* to the total number of time bins in the ATC; *t*_*n*_ is the POI corresponding to the time bin *n*; *ω* is the angular frequency defined as *ω* = 2π*f*, with *f* set to range from 1 to 40 Hz in steps of 0.25 Hz; and *c*_*f*_ is the Fourier coefficient relative to the frequency *f*. This is computed in Matlab using the function “*fft”*.

The complex spectra of all participants were averaged in the complex domain, i.e. taking phase information into account, to obtain a single complex spectrum (equivalent to first averaging the single-participant accuracy time courses over participants in the time domain, and then Hann-tapering, zero-padding and Fourier-transforming them). This complex spectrum was then rectified and squared to obtain the power spectrum (Fig. 3A-F). This method, with slight variation, has been used in several studies (Landau and Fries, 2012).

**Fig. 3.**
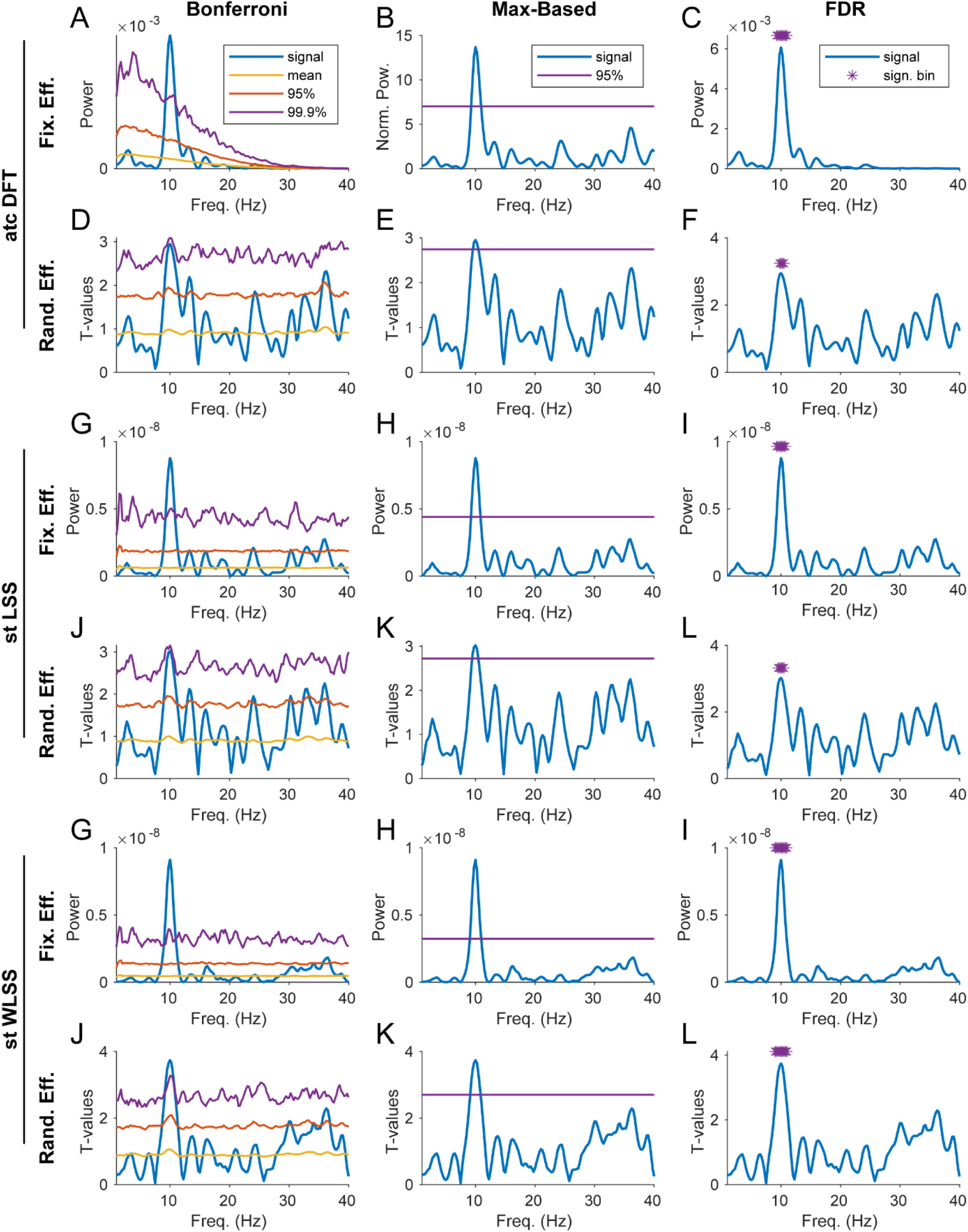
Spectral Analysis. Spectral analysis of the simulated behavioral data generated in Fig. 1, with the calculation of statistical power. The spectral analysis is performed with 3 different methods, atcDFT (A, B, C, D, E, F), stLSS (G, H, I, J, K, L), stWLSS (M, N, O, P, Q, R). The statistical power is calculated for two different test types, a fixed-effect test (A, B, C, G, H, I, M, N, O) and a random-effect test (D, E, F, J, K, L, P, Q, R). The multiple comparison correction is performed in 3 different ways, Bonferroni (A,D,G,J,M,P), Max-Based (B,E,H,K,N,Q), and FDR (C,F,I,L,O,R).

### 2.4. Operating on single-trial data

#### 2.4.1. Preprocessing: tapering and detrending single-trial BRVs

In order to subtract a linear trend from the single trials, we fitted a line to the participant-specific ATC and we subtracted the value of this line at the time point of each trial’s POI from the respective BRV. Similarly, to apply the Hann taper, the value of the taper at the time point of each trial’s POI was multiplied with the respective BRV.

#### 2.4.2. Single-Trial Discrete Fourier Transform (stDFT)

We transformed the single-trial POIs into phases, respectively complex vectors. Each trial’s POI corresponds, for each frequency, to a particular phase, which we call the probe-onset phase *φ* and thereby to a particular complex number. For example, for 10 Hz, with a cycle length of 100 ms, a POI of 50 ms corresponds to 0.5 cycles, which in turn corresponds to a *φ* of π rad, expressed as complex number [−1, 0*i]*. Thus, to represent all frequencies, we use a spectrum of complex numbers. For linearly increasing frequencies, a given POI leads to linearly increasing *φ*, i.e. to a linear slope in the *φ* spectrum.

To obtain the single-trial spectrum, we multiply the *φ* spectrum with the preprocessed BRV of the corresponding trial, separately per frequency. This was repeated for each trial to obtain the cross-spectrum between the alignment event and the (detrended and Hann-tapered) BRVs. We then calculate the single-participant average DFT as the sum of the cross spectra over trials.

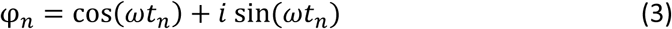

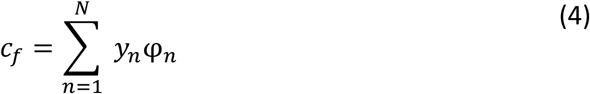

Where *y*_*n*_ is the *n*^th^ element of the vector containing the BRVs, with *n* corresponding to the trial number and *N* to the total number of trials; *t*_*n*_ is the *n*^th^ element of the POI vector; *ω* is the angular frequency defined as *ω* = 2π*f*, with *f* set to range from 1 to 40 Hz in steps of 0.25 Hz; and *c*_*f*_ is the Fourier coefficient for frequency *f*. This can be rewritten as follow:

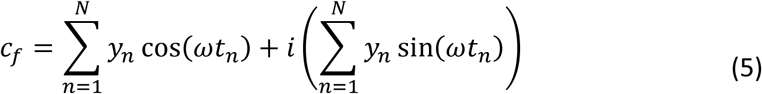

These calculations are done at the level of the single participant, such that each participant gives one complex Fourier spectrum. The complex spectra of all participants were averaged in the complex domain, i.e. taking phase information into account, to obtain a single complex spectrum, which we call the average-participant DFT. This complex spectrum was also rectified and squared.

In the specific case of a regular sampling of the POIs, the stDFT method is equivalent to the atcDFT method, if the latter is performed without applying any convolution (binning being a form of convolution).

#### 2.4.3. Single-Trial Least Square Spectrum (stLSS)

This method consists in calculating a multivariate generalized linear model separately for each participant, using as independent variables, per frequency, the probe onset phases of all trials, and as dependent variable the corresponding BRVs. Note that the Hann-tapering applied to the BRVs allows us to calculate the LSS at any frequency resolution; this is equivalent to Fourier-transformation after zero-padding to the length giving the Rayleigh frequency corresponding to this frequency resolution.

The model behind this analysis can be written as:

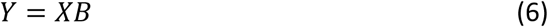

Where:

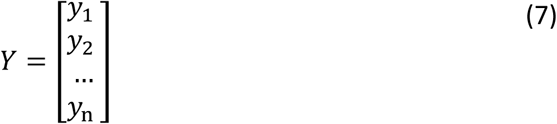

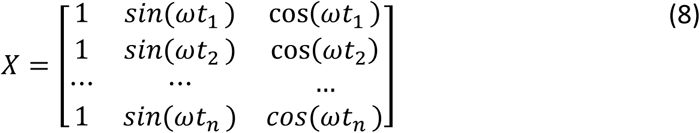

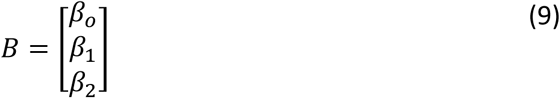

Where *y*_*n*_ is the *n*^th^ element of the vector *Y* containing the BRVs, with *n* corresponding to the trial number and *N* to the total number of trials; *t*_*n*_ is the *n*^th^ element of the vector containing the POIs; *ω* is the angular frequency defined as *ω* = 2π*f*, with *f* set to range from 1 to 40 Hz in steps of 0.25 Hz; and β_*o*_, β_1_ and β_2_ are the regression coefficients.

There is no exact solution to the equation 6, so we preceded with a least square fitting. Solving the minimization problem gives the normal equation and allows us to find the best 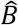 that approximates *B*

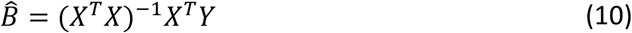

We computed this in Matlab using the backslash operator (as B = X\Y), which is numerically stable.

We therefore obtained the regression coefficients *β*_1_ and *β*_2_, and we used them to calculate the Fourier coefficient relative to the frequency *f*, as follow:

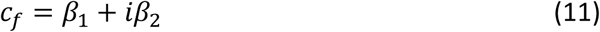

We repeated this procedure for all frequencies of interest so that we had a complex number per frequency, which can be considered as the equivalent of a complex Fourier spectrum. These calculations were done at the level of the single participant, such that each participant gives one complex spectrum. The complex spectra of all participants were then averaged in the complex domain, i.e. taking phase information into account, to obtain a single complex spectrum. This complex spectrum was rectified and squared to obtain the power spectrum (Fig. 3G-L). The stLSS method, with small modifications, has been used in several studies (Fiebelkorn et al., 2013; Tomassini et al., 2017; Benedetto and Morrone, 2019).

Note that the LSS can be applied also to the mean ATC. We refer to it as atcLSS. In this case the *y*_*n*_and *t*_*n*_of equation 7 and 8 have to be defined as follow: *y*_*n*_ is the *n*^th^ element of the vector containing the ATC, with *n* corresponding to the time bin number in the ATC; and *t*_*n*_ is the POI corresponding to the time bin *n*. The preprocessing described for the atcDFT and the negative consequences of convolving the time domain apply here as well. Also here, in the case of a regular sampling, the stLSS method is equivalent to the atcLSS method (when the latter is performed without convolution).

#### 2.4.4. Equivalence of LSS and DFT for regular sampling

The LSS can be considered a more general case of the DFT, able to accommodate irregularly sampled signals. In the specific case of a regular sampling, these two methods are mathematically equivalent. One can observe that the columns of the matrix *X* of equation 8 are orthogonal to each other in the case of a regular sampling (i.e. their dot product is zero). Consequently, *X* is a rectangular semi-orthogonal matrix, and (*X*^*T*^*X*)^−1^ is equivalent to the identity matrix. Therefore, we can simplify equation 10 as follows:

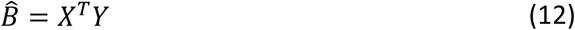

And we can calculate the Beta coefficients as follows:

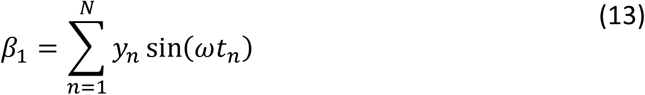

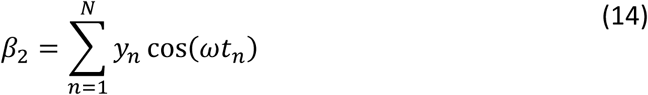

Comparing equation 13 and 14 with equation 2 and 5 shows that in the case of regular sampling, the LSS method is equivalent to the DFT method.

Note, the more the sampling is irregular, the more the stDFT and stLSS will give different results. Yet, for typical levels of sampling irregularity, as simulated here, the results were similar, which suggests that both methods can be equivalently used to estimate the spectrum. Note however, that only the spectrum estimated with the LSS benefits from the time shift invariance property and gives the opportunity to reconstruct the original time domain signal from the frequency domain in a least square sense (VanderPlas, 2018). In this study we simulated the data with irregular sampling, which is more typical of real-world data, and therefore in the figures we illustrate the stLSS method as single-trial method of choice.

#### Single-trial Weighted Least Square Spectrum (stWLSS)

Here we introduce a modified version of the LSS method which takes into account a weighting factor. Factors such as arousal, task engagement and wakefulness may affect the modulation-depth of the observed behavioral rhythms. These factors may be reflected in (or correlated with) parameters such as trial number, pupil diameter, local performances (i.e. performance averaged over the past few trials), heart rate variability, and others. In the simulated data, such an effect was introduced. To take this effect into account in the regression, we calculated the interaction terms between the two sinusoidal components and the vector containing the weights ϒ.

We normalized the weights values in the vector ϒ, so that the sum of these values is equal to the length of the vector. The following procedure for this method is identical to what has been described for the LSS method, except for the matrix *X* in equation 8, which now takes the following form:

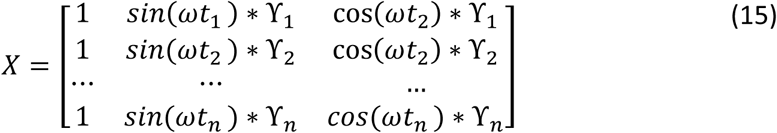

Where ϒ_*n*_ is the *n*^th^ element of the vector ϒ containing the weights; and *ω* is the angular frequency defined as *ω* = 2π*f*, with *f* set to range from 1 to 40 Hz in steps of 0.25 Hz.

### 2.5. Statistics

There are two main questions that we considered with regard to the rhythmicity of behavior: 1) Does behavior show significant rhythmicity, i.e. does rhythmicity exist? 2) Does behavior show different strengths of rhythmicity between two experimental conditions, i.e. does rhythmicity differ?

Furthermore, we can choose to make an inference about the effect on two levels: 1) An inference on the sample of investigated participants, referred to here as fixed-effect analysis (Fig. 3 odd-numbered rows). 2) An inference on the population of all possible participants, referred to here as random-effect analysis (Fig. 3 even-numbered rows). In the case of a random-effect analysis, one might want to weight each participant equally; in this case, the parameter estimates for each participant can be normalized by the number of trials, before combination over participants.

We present non-parametric statistical test methods, because they avoid assumptions about underlying distributions and allow for an elegant way to correct for multiple comparisons across frequencies. The basic approach in these non-parametric statistical test methods is to define a manipulation of the data that would destroy the hypothesized effect (existence or difference), yet not make a difference under the null hypothesis of non-existence or no-difference.

We start by illustrating the random-effect test for a condition difference. In this case, we can use any of the above methods to obtain two spectra per participant, one for each condition. The test statistic quantifies the difference between the two conditions, averaged over participants. This test statistic can be simply the average difference, or a more sophisticated difference metric, like the paired t-test between conditions, across participants. The t-test normalizes the difference by the SD across participants and can thereby equalize e.g. across frequencies. A paired t-test across participants, separately per frequency, gives the observed t-value spectrum. The null hypothesis is that rhythmicity does not differ between the two conditions. Thus, under the null hypothesis, we can randomly exchange conditions, and the resulting t-value spectrum should not change. In the random-effect case, we randomly exchange conditions at the level of the participant. That is: To implement one randomization, we make a random decision, per participant, of whether to exchange the spectra between the two conditions or not. We then proceed as before, arriving at one randomization t-value spectrum. This randomization is repeated many times, thereby giving many randomization t-value spectra. Here, we performed 2000 randomizations. The observed t-value spectrum is then compared to the distribution of randomization t-value spectra. If the observed t-value for a given frequency was smaller than the 2.5^th^ percentile or larger than the 97.5^th^ percentile of the randomization t-value distribution at that frequency, we considered the observed t-value significant with the frequency-wise false positive rate controlled to be below 0.05.

In order to correct for multiple comparisons performed across frequencies there are two main classes of methods. The methods in the first class control the family-wise error rate (FWER), e.g. the probability of at least one false discovery, which is set to be below a critical p-value. The methods in the second class control the false discovery rate (FDR), e.g. the expected proportion of false discoveries over the total number of discoveries, which is set to be below an α-value.

First, we illustrate the Bonferroni correction which belongs to the FWER class. Here the critical p-value is divided by the number of frequencies. For example, for 10 tested frequencies, a p-value of 0.05 would reduce to 0.005, and the relevant percentiles change from the 2.5^th^ to the 0.25^th^ and from the 97.5^th^ to the 99.75^th^ percentile (Fig. 3 left column). This correction is easy to perform, at the cost of sensitivity. Note that the Bonferroni correction should take the true number of independent frequency estimates into account; if zero padding (or effective zero padding in the case of LSS – see above) is used, this increases the number of displayed frequencies in a spectrum, but it does not increase the number of independent frequencies.

An alternative FWER method that comes at less of a cost in sensitivity yet still ensures the required specificity, i.e. that strictly controls the false-positive rate, is the Max-Based correction (Fig. 3 middle column) (Nichols and Holmes, 2002). After each randomization, the maximal t-value across all frequencies is placed into the max-randomization distribution; the minimal t-value across all frequencies is placed into the min-randomization distribution. Note that those randomization distributions lack a frequency dimension. The observed t-value spectrum is compared, frequency per frequency, to those max-and min-randomization distributions. If, for a given frequency, the observed t-value is smaller than the 2.5^th^ percentile of the min-randomization distribution or larger than the 97.5^th^ percentile of the max-randomization distribution, we consider the observed t-value significant with the false positive rate controlled to be below 0.05, corrected for the multiple comparisons across frequencies.

When we apply the Max-Based correction to data obtained using an ATC-based method after convolution in the time domain, we first need to normalize the power spectra, to render the different frequencies comparable. Otherwise, higher frequencies would be attenuated, because of the low-pass filtering effect of the convolution. To this end, we normalized both the observed spectrum and the permuted spectra by dividing by the mean power of all the permuted spectra. We note that this is an ad-hoc solution, which we perform to be able to apply Max-Based corrections to convolution-based spectra at all; this step is not necessary for single-trial based methods, which is an additional point to favor those methods.

Finally, we illustrate the Benjamini-Hochberg procedure for false discovery rate (FDR) correction (Fig. 3 right column) (Benjamini and Hochberg, 1995). Here an *α*-*value* (the expected proportion of false discoveries) has to be chosen. We selected an *α*-*value* of 10%, as it is generally used in the literature (Zhang et al., 2019). Here all the p-values (one for each frequency tested) are ordered from the smallest to largest, and they are ranked. A *critical value* is calculated for each individual p-value as follows:

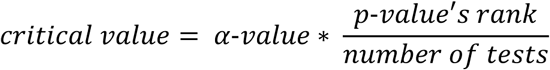

The largest p-value which has a value below its own specific *critical value* is identified, and all the p-values equal to it, or smaller than it, are considered significant.

After illustrating the random-effect test for condition differences, we consider the random-effect test for the existence of a behavioral rhythm. Here, there is only one experimental condition, giving one observed spectrum per participant. One could consider to test against spectra made of zeroes, yet this would ignore the possibility that the observed spectra contain some estimation bias. Therefore, we devised a method to estimate bias spectra per participant. The bias is the value returned by the employed metric in the absence of any rhythmicity. The absence of rhythmicity is equivalent with a situation in which there is no relation between POIs and BRVs. Thus, under the null hypothesis, POIs and BRVs from different trials can be randomly combined, and this should not change the result. We randomly combined POIs and BRVs from the different trials of a given participant to obtain bias estimate spectra. In order to optimize the estimation of the bias estimate spectra, we performed 1000 randomizations per participant, and averaged the spectra to obtain one average bias estimate spectrum per participant. Thus, for each participant, we have one observed spectrum and one average bias estimate spectrum. Further statistical testing, across participants, can compare the observed spectra with the bias spectra, which is similar to the comparison between two experimental conditions described above. One difference in this case is that the test is one-sided, i.e. if the observed t-value is larger than the 95^th^ percentile of the max-randomization distribution, we consider the observed t-value significant.

In some cases, data might be available from only a relatively small number of participants, such that a random-effect test across participants would be very insensitive. In this case, one can consider a fixed-effect test. In the random-effect test, the average difference (either between conditions or between observed and bias spectrum) is compared to the variance across participants. In the fixed-effect test, the same average difference is essentially compared to the variance across the trials pooled over participants. We first consider the case, in which a fixed-effect test compares two conditions. Here, we again combine data from participants e.g. by averaging condition-difference spectra over participants, or by calculating paired t-values spectra across participants. Yet, the randomization proceeds differently, as it has to operate at the trial level. The trials from the two conditions in a given participant are randomly assigned one of the two conditions, such that the number of trials for a given condition remains unchanged (so-called random repartitioning). After this is done for all participants, the condition-wise spectra are calculated per participant, and the spectra are combined over participants as before. This is performed for many randomizations, each time giving one randomization spectrum. Once one observed spectrum and many randomization spectra are obtained, testing proceeds as described above by comparing the observed spectrum with the distribution of randomization spectra.

Finally, we consider the case of a fixed-effect test for the existence of behavioral rhythmicity. In this case, the calculation of the observed spectrum proceeds as for a fixed-effect test of condition differences. Yet, the randomization cannot be based on two conditions, but implements a bias estimate. As in the random-effect case, the bias estimate is based on the random combination of POIs and BRVs of a given participant. In the fixed-effect test, each randomization implements this random combination per participant, calculates a spectrum per participant, and averages those spectra over participants. After many randomizations, this gives many randomization spectra, and the observed spectrum can be compared to the distribution of randomization spectra as explained above.

The fixed-effect testing for the existence of a rhythm is statistically most involved, yet is the situation for most of the relevant studies in the previous literature.

### 2.6. Metrics to quantify the performance of the different methods

To compare the performances of the evaluated methods, we used 3 metrics: Sensitivity, Specificity and D-prime (Fig. 4, 5, 7). In order to calculate these metrics, we generated 300 datasets, changing each time the seed of the random number generator in Matlab, and we applied to each datasets the analysis methods and the statistical approaches illustrated above. Furthermore, we repeated this process for different parameter sets (e.g. we varied the PLV value, the modulation-depth value, the frequency of the underlying rhythm, or the total number of participants).

**Figure 4.**
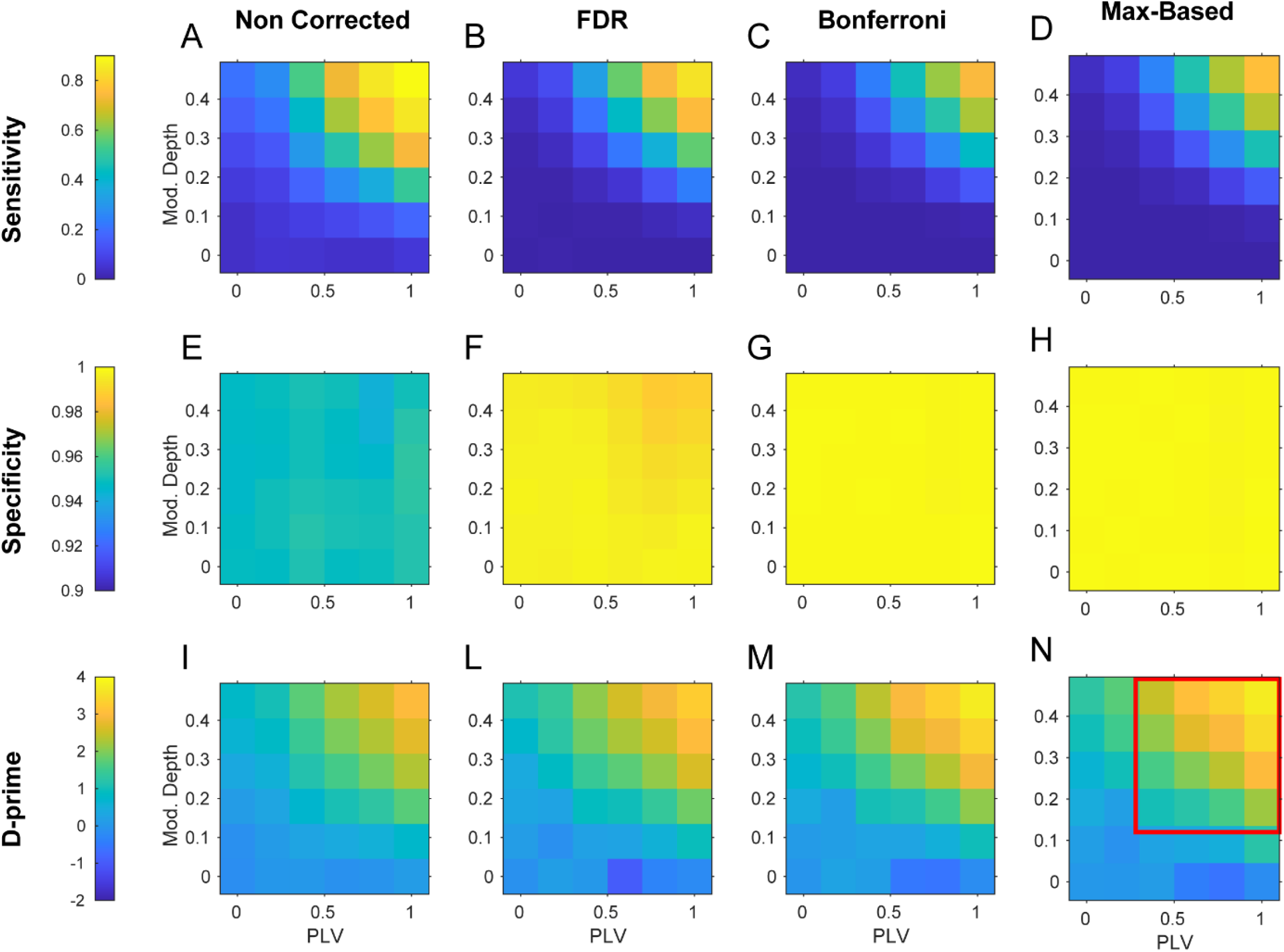
Sensitivity, Specificity and D-prime for variable PLV and Modulation-Depth. Sensitivity, Specificity and D-prime are shown in 2D color plots, for the PLV and Modulation-Depth values indicated on the X- and Y-axes, respectively. The displayed values of Sensitivity, Specificity and D-prime represent the average over 300 simulation runs, in which PLV and Modulation-Depth are set as indicated, and all other parameters are set as in Fig. 1. Different multiple comparison approaches (Non Corrected, FDR, Bonferroni and Max-Based) are applied in the four columns.

**Figure 5.**
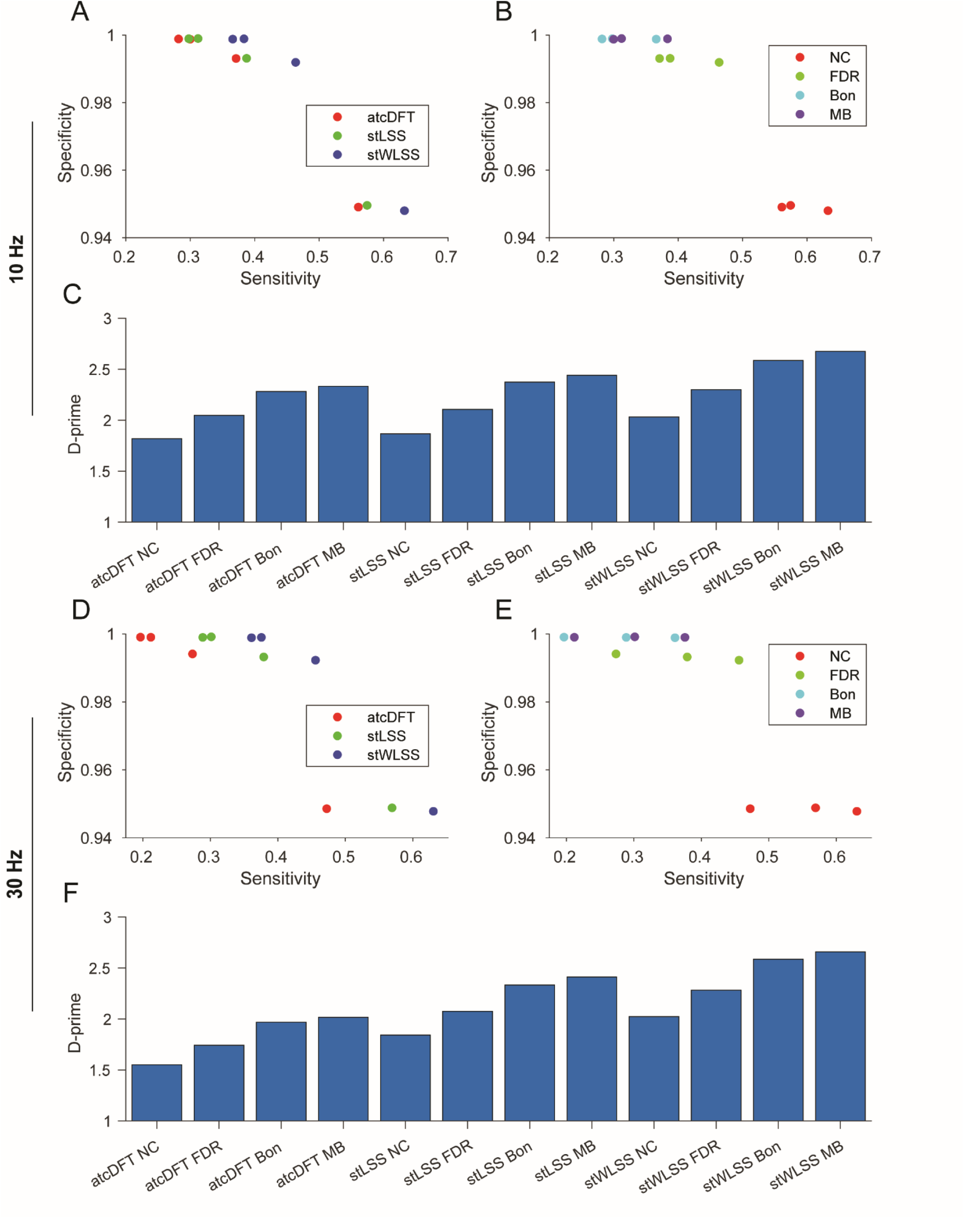
Comparison of methods’ Performances. (A,B,C) We compare Sensitivity, Specificity and D-prime for the values generated from averaging the results of Fig. 4 in the range of PLV and Modulation-Depth values indicated by the red rectangle in Fig. 4N. Sensitivity and Specificity are visualized in two scatter plot, where the dots are colored according to the analysis method (A), and according to the multiple comparison correction approach (B). The corresponding D-prime values are visualized as a bar graph (C). (D,E,F) Same as in the previous panels, for data generated with identical parameters, except for the underlying rhythmicity which in this case was set to 30Hz

The applied methods and statistical approaches are computed for many frequency bins, including frequency bins corresponding to spectral interpolation. We need to determine for which frequency bins a significant result is considered a *Hit*. When we simulate an underlying 10 Hz rhythm in our dataset, we expect the applied methods to report significant rhythmicity for the frequency bins at and around 10 Hz, and not for frequency bins far from 10 Hz. We decided to consider as *Hit*s all the frequency bins which are included between the simulated frequency +/- 1.5 Hz, and which are reported as significant. We chose this frequency range because it corresponds to the clusters of significant frequency bins that we obtain for the simulated datasets with higher PLV and higher modulation-depth. When one frequency bin in this range was identified as non-significant, it was considered a *Miss*; when a frequency bin outside this range was identified as significant, it was considered a *False Alarm*; and when a frequency bin outside this range was identified as non-significant, it was considered a *Correct Rejection*.

We defined Sensitivity, Specificity and D-prime as follows:

#### Sensitivity

We defined Sensitivity as the hit rate:

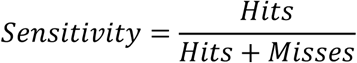

#### Specificity

We defined Specificity as the correct rejection rate:

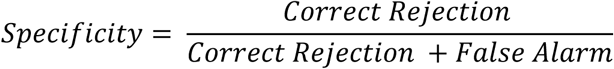

#### D-prime

To calculate the D-prime, the general formula is:

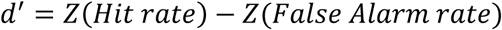

Where the function *Z*(*x*) is the inverse of the cumulative distribution function of the Gaussian distribution. Because both *Hit rate* and *False Alarm rate* in our simulation can take values of 0 and 1 and the function *Z*(*x*) for these values is equal to +/- infinite, we applied the log-linear rule (Hautus, 1995). Accordingly to this correction, the *Hit rate* and the *False Alarm rate* were calculated as follow:

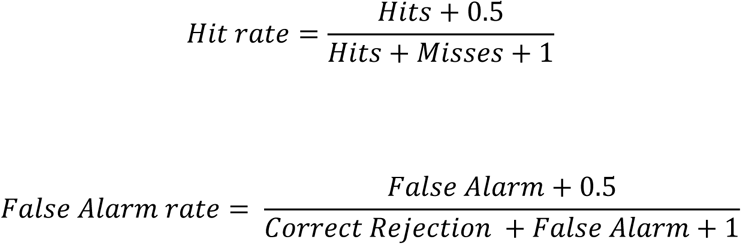

### 2.7. Exploring PLV and Modulation-Depth parameter space

We ran sets of 300 simulations, for different combinations of modulation-depth and cross-participant PLV values (as specified in Fig. 4, x- and y-axes), each simulation containing 30 participants with 400 trials each. For each set, the metrics described above were calculated. This was separately performed simulating an underlying rhythmicity both at 10 Hz and at 30 Hz.

In Figure 4 we visualize the pattern of results obtained with the stDFT method, a fixed-effect statistical approach, a simulated rhythm at 10 Hz, and different multiple corrections. The patterns obtained with the atcDFT and the stWLSS methods are not shown, because they are qualitatively very similar, and a comparison between fixed-effect and random-effect is addressed later (Fig. 7).

In Figure 5 we compared Specificity, Sensitivity, and D-prime for different analysis methods combined with different multiple comparison corrections approaches, both for datasets with a simulated 10 Hz (Fig. 5A-C) and 30 Hz (Fig. 5D-F) rhythm. The reported values are the results of an average of these values obtained for simulation sets with different PLV and modulation-depth values (in the range indicated by the red rectangle of Figure 4, panel N).

The single-trial methods exhibit higher Sensitivity and D-prime than ATC-based methods. This difference notably increases for higher frequency rhythms. The stWLSS method, taking into account an extra weighting factor, further increases Sensitivity and D-prime in respect to the stLSS. The Max-Based and Bonferroni corrections exhibit lower Sensitivity but higher Specificity than the FDR correction. When Sensitivity and Specificity are combined to give the D-prime, the Max-Based correction performs best.

### 2.8. Phase-aligned within subjects (PAWS)

So far, we assumed that there is some phase alignment across trials within a participant, and also some phase alignment across participants. We refer to this as “Phase-Aligned Within And Across Subjects” or PAWAAS. This assumption renders our analysis more sensitive, because the quantification can minimize the influence of random fluctuations with random phase across participants. This is accomplished by the averaging of complex spectra over participants, which corresponds to an averaging in the time domain, which in turn leads to partial cancellation of any fluctuations that are not phase aligned across participants. Yet, in some cases, an investigator might not want to subscribe to this assumption, and rather test whether there is a rhythm, irrespective of phase alignment across participants. This can be accomplished using almost exactly the same methods as described above, with a single modification. For the quantification of the phase-aligned rhythms, we averaged the complex spectra over subjects and then took the absolute magnitude; for the quantification of the non-phase-aligned rhythms, we took the absolute magnitude of the complex spectrum of each participant, and then calculated the average over participants. All other steps remained the same. As this approach requires phase alignment only within subjects, we refer to it as “Phase-Aligned Within Subjects” or PAWS. An illustration of the consequences of the PAWAAS versus the PAWS assumption for cases of high (0.92) and lower (0.4) PLV using stLSS are shown in Fig. 6.

**Fig. 6.**
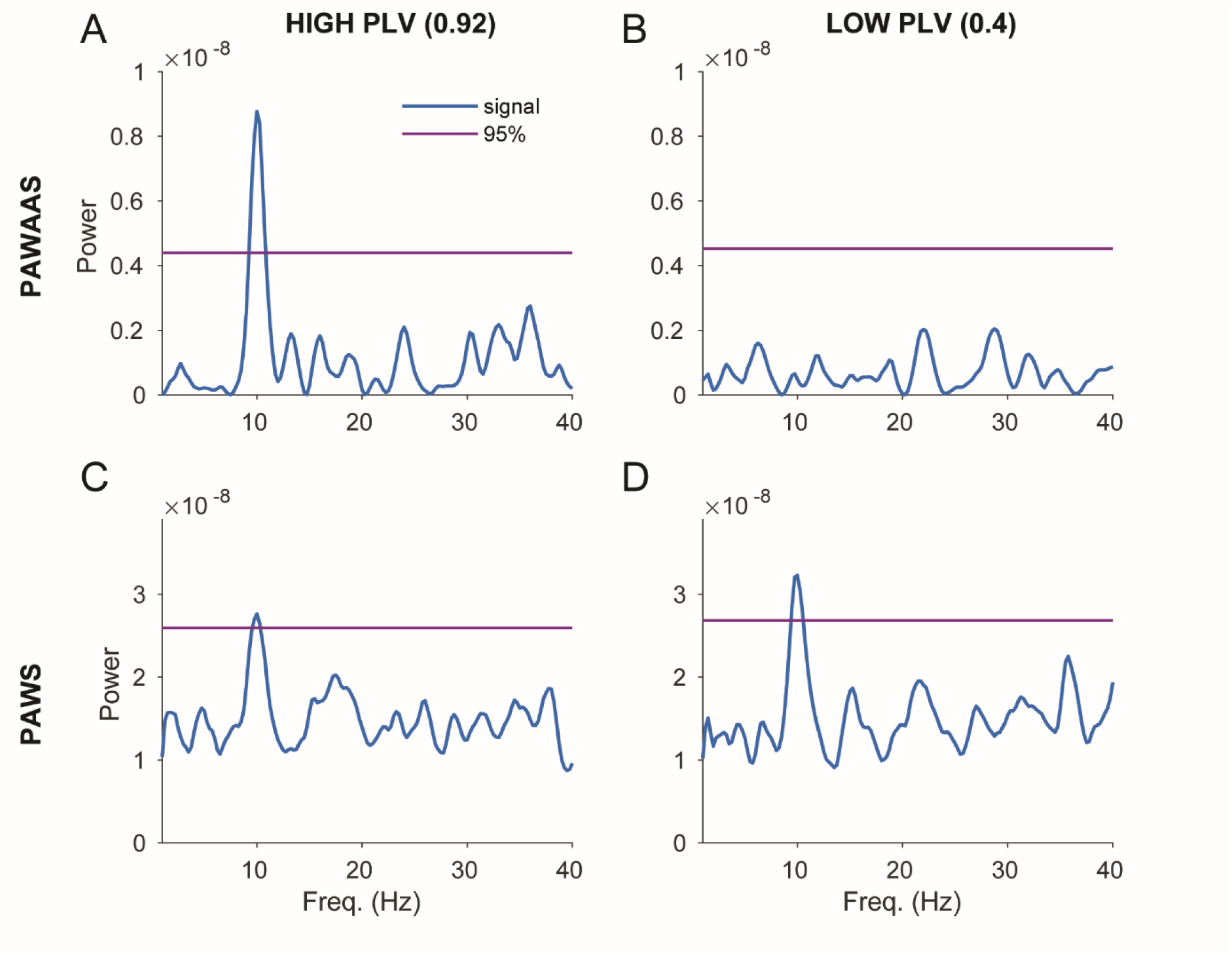
PAWS and PAWAAS. Illustration of the different application of PAWS and PAWAAS using the stLSS method. The two approaches are applied to the same data generated in Fig 1 (A, C) and to data generated with a lower PLV across participants (0.4 instead of 0.92) and otherwise identical parameters (B, D). PAAS performs better than PAWAAS when the PLV is high (A, C), while PAWAAS performs better when the PLV is low (B, D).

### 2.9. Dampened Oscillator

The simulated average accuracy time course resembled a dampened harmonic oscillation (Fig. 1C). This is due to the fact that after the alignment event, the rhythm’s phase is maximally consistent across trials and/or participants, while it later decorrelates. Furthermore, averaging over trials and participants with slightly different modulation phase and frequency also leads to dampening. We therefore explored the possibility to fit the accuracy time course with a dampened oscillator (Fig. 2C, D). This has already been done in the rhythmic-sampling literature by a study evaluating rhythmicity in the decoding accuracy of MEG data (Wutz et al., 2016). We fitted the following function to the mean accuracy time course:

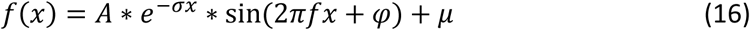

where *A, σ f, φand* µare free parameters representing respectively the amplitude, the exponent of the exponential component, the frequency and the phase of the sinusoidal component, and an offset. To find the parameters which are minimizing the squared error, we used the Matlab function “*lsqcurvefit*”. When multiple models are fit to the same time series, the Akaike Information Criterion (AIC) can be used to provide a means for model selection.

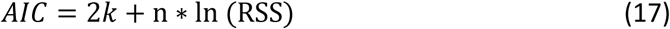

Where *k* is the total number of estimated parameters; n is the number of time points; and RSS is the “residual sum of squares” of the fitting. The fit of a dampened harmonic oscillation in our simulations always provided a lower AIC than the fit of a sine.

This could be applied also to the DFT based methods using dampened sinusoids as basis functions and to the LSS based methods using dampened sinusoids as regressors. In this case, one would have to explore a range of values for the exponent of the exponential component, and to choose the one which minimizes the squared error. An extensive illustration of this implementation is beyond the scope of the current study.

### 2.10. Optimal division of a fixed total number of trials across a selectable number of participants

A practical decision when recording behavioral performance data is whether it is more advantageous to collect more trials per participant, or fewer trials from more participants. The trade-offs here are not necessarily obvious, and may depend on each experimenter’s desired level of inference. To find the optimal distribution of trials across subjects, we fixed the total number of trials at 32000, and distributed those trials equally over a variable number of participants, namely either 8, 16, 32, 64, or 128 participants, both in the case of a simulated rhythmicity at 10 Hz and at 30 Hz. For each condition we ran a set of 400 simulations, and we calculated Specificity Sensitivity and D-prime for the different methods, and for the different statistical approaches (random-effect and the fixed-effect).

In Figure 7 we illustrate the results. For single-trials methods performed with a random-effect statistical approach distributing the same number of trials among a larger number of participants benefits Sensitivity and D-prime. Note that this is not due to our simulation containing less variability across participants than across trials, because these parameters were matched. The same manipulation for single-trials methods performed with a fixed-effect statistical approach does not affect Sensitivity and D-prime. A very different trend is observed for ATC-based methods. Here distributing the same number of trials among a larger number of participants has a detrimental effect for Sensitivity and D-prime, in particular when the simulated rhythm is of higher frequency. This because having less trials per participant corresponds to a sparser sampling of the POI time, and a convolution in the time domain of such a signal introduces larger distortions in the frequency domain, in particular for higher frequencies.

**Fig. 7.**
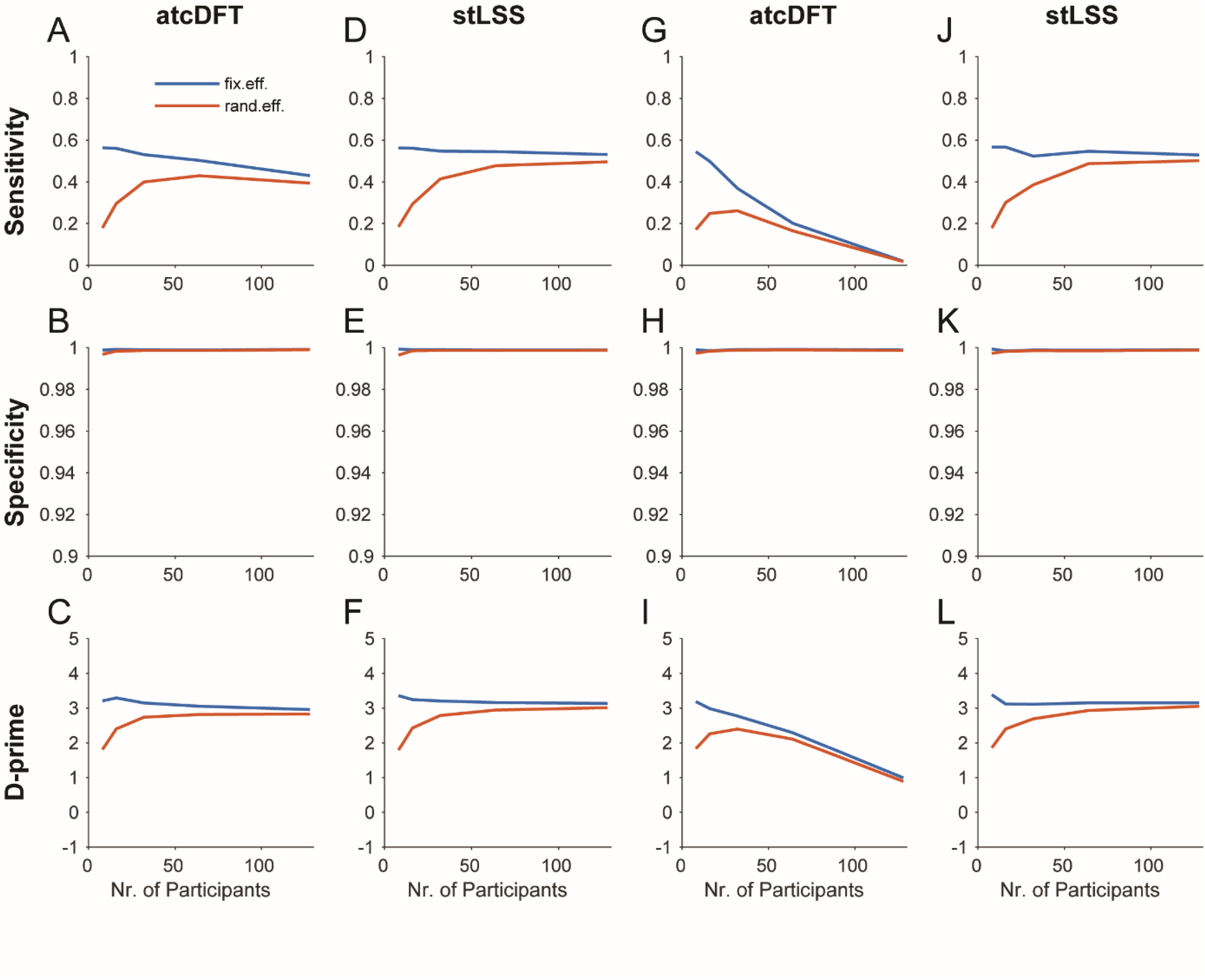
Varying participant number, keeping the total number of trials constant. We show how Sensitivity, Specificity and D-prime values change, when the number of participants is varied and the total number of trials is kept constant. The results are shown both for a fixed effect (blue line) and a random effect (red line) statistical approach, for the atcDFT (even columns) and the stLSS (uneven columns) methods, and for data simulated with an underlying rhythmicity of 10 Hz (A-F) and of 30 Hz (G-L). The data are generated keeping the total number of trials constant to 12800, and varying the total number of participants with the following values: 8, 16, 32, 64, and 128. All other parameters were set as in Fig 1.

Note that Sensitivity and D-prime cannot be directly compared between fixed-effect and random-effect tests, because those two types of tests provide qualitatively different inferences: the fixed-effect test provides an inference on the investigated sample of participants, whereas the random-effect test provides an inference on the population from which the investigated participants have been sampled.

Figure 7 shows the pattern of results obtained with the stLSS method and a Max-Based multiple comparison correction. The patterns obtained with the atcDFT and the stWLSS methods are not shown, because they are qualitatively very similar. The patterns obtained with other multiple comparison corrections shows the same differences already illustrated in Figure 4 and 5.

## 3. Discussion

The number of studies investigating rhythms in behavior has been growing in recent years (VanRullen, 2016a). However, different methods have been used to analyses these data, making the comparison between studies difficult, and concerns over reproducibility have been raised (Lin et al., 2021; Sun et al., 2021; van der Werf et al., 2021). Moreover, it is not clear which of these methods is more optimal to maximize the detection of true rhythms while minimizing the chance of false discoveries. In this paper, we simulated ground-truth data, using a model resembling a typical experiment. We then analyzed the ground-truth data with several methods, and compared their performance using Sensitivity, Specificity and D-prime.

We identified two main classes of methods for spectral analysis: ATC-based methods (atcDFT, atcLSS), and single-trial methods (stDFT, stLSS). In the case of a regular sampling, these methods are analytically equivalent, but not in the case of an irregular sampling, which is more typical of real-world data. In the case of an irregular sampling, the methods based on time-averaged data require as additional pre-processing step a convolution in the time domain, and this introduces the following disadvantages. First, the convolution has a low-pass filtering effect, which precludes a direct interpretation of the spectra with regard to the presence of a rhythm and it reduces the sensitivity for the detection of rhythmicity at higher frequencies. Second, the low-pass filtering effect has non-beneficial consequences for statistical testing, requiring an additional normalization for the Max-Based approach. Third, the convolution in the ATC-based methods weighs each time point equivalently (even when different time points are the result of an average over different numbers of trials), while single-trial methods give each trial the same weight. In terms of Sensitivity, Specificity and D-prime, we found that the single-trial methods performed better. The differences between the two classes of methods grow as the rhythm moves to a higher frequency and as the sampling becomes more irregular.

There are two main single-trial methods, namely stDFT and stLSS. Even if stLSS and stDFT are giving similar results when applied to our simulated data, the stLSS benefits of additional mathematical properties which makes it more appropriate when dealing with very sparsely sampled signals. Additionally, the stLSS can be easily modified to stWLSS, offering the possibility to include a weighting factor like the pupil size on a given trial or the behavioral performance in the recent history of trials, which can further improve performance.

For the analysis of experimental data, we suggest the stLSS method as the first choice. The stLSS method is appropriate for both regularly and irregularly sampled data. If an additional modulatory factor, possibly influencing the strength of the underlying rhythm, is measured, we recommend the stWLSS method.

In recent years most of the rhythms reported in behavioral data were in the theta (Landau and Fries, 2012; Holcombe and Chen, 2013; Tomassini et al., 2015; Hogendoorn, 2016; Senoussi et al., 2019; Benedetto et al., 2021; Plöchl et al., 2021), alpha (Song et al., 2014; Benedetto and Morrone, 2019; Ho et al., 2019; de Graaf et al., 2020) and beta (Bell et al., 2020; Veniero et al., 2021) range. However these studies primarily employed ATC-based methods, which are suboptimal to detect rhythms of higher frequency. Therefore, it is possible that the relative lack of evidence for higher-frequency behavioral rhythms, such those in the gamma range (30-100Hz), may be due to the methods rather than the lack of behavioral rhythms in this frequency range (but see (Dehaene, 1993)).

A particular ATC-based method used in the literature is sine fitting. This method requires a convolution in the time domain for irregularly sampled signals, which brings all of the previously mentioned shortcomings. Moreover, sine fitting does not provide a full characterization of the spectra, which would be useful to infer the presence of an underlying periodic process. However, there are some useful applications for this method. For example, the experimenter can explore the possibility of fitting a damped harmonic oscillator instead of a simple sine wave, which may be more appropriate to represent time-averaged data after a reset event. Theoretically speaking, this ATC-based method allows the possibility to fit any arbitrary modulation function to the data, and a diverse set of modulations can be tested simultaneously. When different models are fit to the same time-series, the best model can be chosen accordingly to the Akaike Information Criterion (AIC), which aims to maximize the trade-off between the goodness of fit and the complexity of the model (Parzen, 1998; Burnham and Anderson, 2004).

We also presented a method that is sensitive to phase alignment within but not across subjects, which we called PAWS. Such a constellation of phase alignments has, to the best of our knowledge, not been reported so far, yet it might exist, and the presented methods can test for it. Note that if there is some degree of phase alignment across participants, then testing for phase-aligned rhythmicity using PAWAAS will increase the Sensitivity, because this method cancels random variability within individual participants that contributes power per participant, but not in the time or phase average over participants.

In this study, we illustrated the case of rhythmicity detection, addressing the question whether there is more rhythmicity than expected by chance, while in some cases the experimenter might aim for rhythmicity comparison, addressing the question whether rhythmicity differs significantly between two conditions. In the latter case, the two conditions may have a different number of trials. This case can often arises when conditions are formed posthoc, e.g. comparing correct versus incorrect trials and performance is not exactly 50%. In this context, it is relevant to keep in mind that differences in sample size might incur different bias values in the employed metrics. To avoid those biases, the condition with the larger number of trials should be subsampled, such that the metrics is repeatedly applied to samples of the same size as the condition with the smaller number of trials, and subsequently, the subsample-based metrics are averaged.

We compared different ways of performing multiple comparison corrections for the statistical analysis, namely False Discovery Rate, Bonferroni, and Max-Based. False Discovery Rate has higher Sensitivity but lower Specificity, while Max-Based and Bonferroni have a lower Sensitivity and a higher Specificity. When combining Sensitivity and Specificity in the D-prime metric, False Discovery Rate performs poorly, followed by Bonferroni, and Max-Based is the best performing method. We suggest to consider False Discovery Rate only in pilot phases of a study, yet to use the robust control of the false-positive rate provided by Max-Based for the final results.

We illustrated two different statistical approaches, a random-effect and a fixed-effect test. A random-effect test allows for an inference on the population, while a fixed-effect test allows an inference on the investigated sample of participants (Fries and Maris, 2021; Combrisson et al., 2022). We therefore suggest to aim for a random-effect analysis whenever possible. We further explore how the two approaches perform, varying the number of simulated participants, while keeping the total trial number constant. We show that when a single-trials method is applied to the data with a fixed-effect test, Sensitivity and D-prime are only slightly affected by how we distribute our trials across participants. Conversely, when a random-effect test is used in combination with a single-trial method, Sensitivity and D-prime benefit from a larger number of participants, even if the number of trials per participant is correspondingly lower. Note also that in practice, each participant requires substantial time to recruit and to set up (and potentially to train in the task), such that a decent number of trials, e.g. what can be achieved within a comfortable session time, appears as a good solution. However, our simulations suggest that a putative approach of bringing a given participant back for multiple sessions is less effective than using the further sessions for further participants.

Note that the analysis methods presented here in the context of a detection task could apply in a similar way also to the behavioral responses in a discrimination task, or to the analysis of reaction times. When the experimenter choose a discrimination task to investigate rhythmicity after a reset, there is the additional opportunity to separately analyze sensitivity and response bias (Ho et al., 2017; Benedetto and Morrone, 2019). This can be done also using a single trial method (Ho et al., 2019).

It is possible to adopt a slightly different experimental approach to the one modeled in our simulation to investigate similar questions: Instead of using an aligning event such as a motor action or an external flash, the time of the probe event can be transformed into a spectrum of phases relative to brain activity recordings, e.g. with EEG or MEG. This experimental approach allows to trace a more direct link between behavioral rhythms and brain rhythms. With regard to the data analysis, in this case there will also be one spectrum for each trial, and these spectra have to be combined over trials. At this point, a very similar logic to the one illustrated above might be applied, and a similar comparison as presented here would be informative (VanRullen, 2016b; Zoefel et al., 2019; Lundqvist and Wutz, 2022). Additionally, these spectra also contain meaningful amplitude information, allowing to weight each trial by its spectral amplitudes.

A recent preprint (Brookshire, 2021) raised the point that testing the observed data against randomization distributions obtained by randomly pairing BRVs and POIs from different trials allows to reject the null hypothesis that there is no consistent temporal structure in the accuracy time course. However, Brookshire (2021) claims that this does not allow to distinguish between periodic and aperiodic temporal structures. We argue that a correct discrimination between periodic and aperiodic temporal structures can be based on a parametrization of the spectrum. How to parametrize a spectrum and how to then quantify the degree of periodicity is beyond the scope of this paper, but it is currently under discussion in the field of electrophysiology (Donoghue et al., 2020; Gerster et al., 2022).

Here we want to clarify that the significance of the test at a frequency bin *f*, and the conclusion that the reset is followed by a significant phase-locked behavioral modulation at *f* is valid independent of whether the underlying process is periodic. Randomly pairing BRVs and POIs from different trials is equivalent to randomly pairing BRVs and probe onset phases from different trials. Thereby, the null hypothesis that behavioral performance is not dependent on POI, is equivalent to the null-hypothesis that performance is not dependent on frequency-wise probe-onset phase. Thus, if statistical tests are significant for a given frequency bin, they demonstrate that for this frequency bin, behavioral performance significantly depends on that frequency’s probe-onset phase.

However, a significant frequency bin *f* in isolation can be either an indication of phase alignment of a periodic process at that frequency, or it can be part of a spectral pattern characteristic of an aperiodic process, e.g. of a 1/ *f* pattern, or of a pattern that is entirely flat across frequencies. To move from a spectrum to the inference on a likely underlying periodic or non-periodic process, one has to consider the entire spectrum or at least a substantial part of the spectrum. Moreover, for a correct interpretation it is crucial to calculate a spectrum without introducing any distortions in the processing steps. We discourage the use of methods operating on trial-averaged data, because they often require a convolution in the time domain which acts as a low-pass filter, reducing the power of high frequencies. Furthermore, we discourage the use of polynomial detrending of order higher than one, because this can act as a high-pass filter, reducing the power of low frequency. These filters, in particular when combined, can produce a spectral shape suggestive of a periodic process, where in fact the original data did not contain such periodicity. Finally we caution against spectral peak alignment across subjects, as this could result in a final spectrum giving a strong impression of rhythmicity even when the data contains only noise (van der Werf et al., 2021).

Generally, the framework presented here, and the provided code, can be used to quantify the performance of any novel metric and a quantitative comparison to the existing metrics.

## Abbreviations

AIC: Akaike Information Criterion
ATC: Accuracy Time Course
atcDFT: Discrete Fourier Transform applied to the mean accuracy time course
atcLSS: Lest Square Spectrum applied to the mean accuracy time course
BRV: Behavioral Response Values
DFT: Discrete Fourier Transform
EEG: electroencephalography
FDR: False Discovery Rate
FWER: Family-Wise Error Rate
Hz: Hertz
LSS: Lest Square Spectrum
MEG: magnetoencephalography
ms: milliseconds
PAWAAS: Phase Alignment Within And Across Subjects
PAWS: Phase Alignment Within Subjects
POI: Probe Onset Interval
RSS: Residual Sum of Squares
SOA: Stimulus-Onset Asynchrony
stDFT: Discrete Fourier Transform applied to single trials
stLSS: Lest Square Spectrum applied to single trials
stWLSS: Weighted Least Square Spectrum applied to single trials
TMS: Transcranial Magnetic Stimulation

## 4. Acknowledgements

We thank Alessandro Benedetto for the conversations on how he applied the sine fitting and the LSS methods in his studies.

## 6. Author contributions

**Tommaso Tosato:** Conceptualization, Methodology, Software, Formal analysis, Writing – Original Draft, Writing – Review & Editing, Visualization. **Gustavo Rohenkohl:** Conceptualization, Methodology, Writing – Original Draft, Writing – Review & Editing, Supervision, Funding acquisition. **Jarrod Robert Dowdall:** Conceptualization, Methodology, Software, Writing – Review & Editing. ***Pascal Fries:*** Conceptualization, Methodology, Writing – Original Draft, Writing – Review & Editing, Supervision, Project administration, Funding acquisition.

